# Co-dissemination of anti-phage defence systems and multidrug resistance is associated with the seventh-pandemic *Vibrio cholerae* defensome

**DOI:** 10.64898/2026.07.13.738241

**Authors:** M. Mozammel Hoque, Joyce To, Dominic Leo, Diane McDougald, Jyl S. Matson, Gustavo Espinoza-Vergara

## Abstract

Anti-phage defence systems in *Vibrio cholerae* are increasingly recognised as cargo on mobile genetic elements (MGE) that also carry antimicrobial resistance (AMR) and virulence genes, but a population-scale view of how the *V. cholerae* defensome partitions between the pandemic O1 clade and the environmental non-O1 reservoir has been missing. We analysed 1,740 RefSeq *V. cholerae* genomes (773 O1, 967 non-O1) sampled from 1934 to 2022 across 60 countries. The non-O1 reservoir harboured 283 defence subtypes and 55 families absent from any O1 genome. O1 was dominated by abortive infection systems (33.5% vs 14.0% in non-O1; P < 0.001). Defence and prophage burden were inversely related within O1 (R = −0.15) but weakly positive in non-O1, and an 80-year reconstruction revealed mirror trajectories of rising defence diversity and declining prophage burden in O1, coincident with the emergence of SXT/R391 ICE-borne AMR. A bipartite network identified strong defence-AMR associations (Phi up to 0.80), nominating AbiE, AbiJ, dCTPdeaminase, Menshen, Retron-I_A and Lamassu-Fam as candidate co-acquired cargo. UMAP clustering resolved seven distinct defensome lineages, and a Random Forest classifier trained on the defence profile alone predicted-MDR (genotypic MDR) status with pooled cross-validated AUC of 0.96, AUC of 0.92 within O1 only and 0.93 within non-O1 only and generalised across decades (AUC = 0.88) and regions (AUC = 0.92); a five-system signature (AbiE, BREX-I, CBASS-II, dCTPdeaminase, AbiJ) achieved AUC = 0.91. These data implicate the AMR-bearing mobilome rather than phage predation as the principal driver of anti-phage defence evolution in pandemic *V. cholerae*, and support further evaluation of the defensome as a genomic correlate of multidrug resistance for surveillance use, pending validation in prospectively phenotyped cohorts.

**Impact statement:** The seventh cholera pandemic, ongoing for more than six decades, is driven by a single *Vibrio cholerae* lineage whose success has been linked to a small set of horizontally acquired mobile genetic elements. Recent work has shown that these elements also carry anti-phage defence systems, but the population-scale picture has been bounded by small genome sets and short clinical sampling windows. We provide a pan-genomic map spanning 1934 to 2022, based on annotation of 1,740 publicly available *V. cholerae* genomes (with quantitative reconstruction limited to 1970 onwards, where per-window sample sizes support it). We identify the non-O1 environmental reservoir as a substantially larger source of bacterial immune diversity than previously appreciated, document a qualitatively different defence-mechanism balance between pandemic and environmental compartments, and nominate specific candidate defence systems for future mechanistic and long-read characterisation as co-occurring cargo on the SXT/R391 family of integrative conjugative elements. The resource and findings provide an evidence base for *V. cholerae* genomic surveillance and inform the design of phage-therapy efforts directed at circulating pandemic lineages.

## Introduction

Cholera remains a major cause of acute diarrhoeal morbidity and mortality worldwide, with an estimated 1.3 to 4 million cases annually [1]. The ongoing seventh pandemic, which began in 1961, is driven almost exclusively by the toxigenic O1 El Tor lineage of *Vibrio cholerae* (seventh pandemic El Tor; 7PET) [2, 3]. Although *V. cholerae* comprises more than 200 serogroups, only O1 and O139 have ever caused pandemic disease, and the emergence of 7PET has been mapped to a small set of horizontally acquired mobile genetic elements (MGEs) including the CTXΦ prophage, the *Vibrio* pathogenicity islands VPI-1 and VPI-2, the *Vibrio* seventh pandemic islands VSP-I and VSP-II, and the SXT/R391 family of integrative conjugative elements (ICEs) carrying the canonical multidrug-resistance cassette *dfrA1*, *sul2*, *floR*, *aph(6)-Id*, *aph(3’’)-Ib* and *catB9* [4–7].

Over the past five years, the long-standing virulence- and AMR-centered view of these MGEs has been substantially extended. Two plasmid defence systems, DdmABC and DdmDE, were identified on VSP-II and VPI-2 respectively in pandemic *V. cholerae* [8], and structural work subsequently resolved DdmDE as a DNA-guided Argonaute-helicase complex that selectively targets small multicopy plasmids [9, 10]. The VPI-2 island was further shown to encode two distinct restriction-modification systems, T1RM and TgvAB [11], while the West African-South American pandemic lineage carries an additional defensive repertoire WonAB, GrwAB and VcSduA on the WASA-1 prophage and a lineage-specific VSP-II variant [12]. Beyond the core genomic islands, phage-inducible chromosomal island-like elements (PLEs) were recognised as widespread phage-defence loci across 7PET strains [13], and the chromosomal integron has more recently been reframed as a biobank of anti-phage defence cassettes available for horizontal mobilisation [14]. Classical biotype *V. cholerae* additionally harbours a SIR2-like NAD -depleting defence system [15]. Taken together, these findings have consolidated a conceptual shift that the evolutionary success of pandemic *V. cholerae* is shaped not only by classical virulence factors but also by a rich and expanding arsenal of anti-MGE immunity [16, 17].

A crucial mechanistic insight came from 34 months of longitudinal clinical sampling in Bangladesh, which demonstrated that SXT/R391 ICEs encode anti-phage defence within a single hotspot of genetic exchange, and that phage predation actively stimulates SXT ICE conjugation, driving concurrent co-transfer of defence and antibiotic resistance to recipient cells [18]. This finding established the principle that defence and AMR can travel together on the same mobile element, but it was necessarily bounded by a short sampling window and a focus on a single genomic hotspot. Whether the same co-dissemination signature is detectable at population scale, whether it extends to defence systems beyond the BREX and restriction-modification hotspot already characterised within the SXT ICE, and how the pattern has evolved across the eight decades of the seventh pandemic have all remained open questions. The most directly relevant pan-genomic perspective to date comes from an analysis of the mobilome and defensome of 46 *V. cholerae* strains, which explicitly proposed that non-pandemic lineages may serve as reservoirs for emerging defence strategies while noting that the genetic diversity of the non-pandemic compartment remains poorly characterized [19]. Sustained improvements in defence-system detection, in particular the DefenseFinder pipeline and its underlying Macromolecular System Finder (MacSyFinder) framework [20, 21], now make a substantially expanded pan-genomic analysis tractable.

Here we address three specific gaps. First, the size and composition of the environmental *V. cholerae* defence pangenome, and how much of it is shared with the pandemic clade, remain uncharacterised at population scale. Second, the temporal dynamics of pandemic defence content over the eight decades of the seventh pandemic, and how these relate to prophage carriage, plasmid acquisition and AMR emergence, have not been systematically mapped. Third, the identity of defence systems co-acquired with the SXT/R391 AMR cassette beyond the known BREX/RM hotspot and the VPI-2/VSP-I/VSP-II cargo has not been resolved. We address these gaps by integrative analysis of 1,740 *V. cholerae* RefSeq genomes (773 O1 and 967 non-O1) sampled across 60 countries between 1934 and 2022.

## Methods

### Genome collection and curation

We assembled a cohort of 1,740 *V. cholerae* genomes available from the NCBI RefSeq database as of January 2022. For a small subset of isolates with deposited raw reads but no precomputed assembly, paired-end Illumina reads were retrieved with the NCBI SRA Toolkit (fasterq-dump v2.11.0) and assembled de novo with SPAdes v3.11.1 under default parameters [22]. All downloaded and locally assembled genomes were then processed through the PanACoTA v1.4 prepare and annotate modules with default filtering threshold, which inherently apply per-genome quality control [23]. Sample-level metadata (country, isolation year, source, host) were retrieved with the NCBI EDirect utilities (esearch, efetch, xtract) queried against the BioSample and SRA databases, and free-text fields were normalised with custom AWK scripts [24]. Countries were aggregated into seven epidemiological regions (South Asia, Africa, Caribbean and Latin America, Europe, Southeast Asia, North America, East Asia) following global cholera surveillance conventions [2, 3]. Serogroup status was assigned in silico by BLASTn (blastn v2.13, -evalue 1e-20, -perc_identity 90, -qcov_hsp_perc 80) of the canonical *ctxB* and rfb O1 gene-cluster references (NCBI accessions AE003852 *ctxAB* and X59554 rfb-O1 wbe operon) against each assembled genome; isolates with a positive *rfb*-O1 hit were classified as O1 and the remainder as non-O1, and *ctxB* presence was scored separately as a toxigenicity marker. To ensure a consistent annotation basis for downstream analyses, all assemblies were reformatted to gembase nomenclature with PanACoTA v1.4 [23], which renames contigs as VICH.1022.NNNNN and produces standardised protein FASTA, GFF and LSTINFO files; gene calling was performed by Prokka v1.14 [25] embedded within the PanACoTA pipeline.

### Anti-phage defence and mobile element detection

With curated assemblies and standardised annotations, we applied a suite of orthogonal genome-mining tools to catalogue four content classes per isolate: anti-phage defence systems, antimicrobial resistance genes, virulence factors and mobile genetic elements. Anti-phage defence systems were detected with DefenseFinder v1.2.0 [20], which runs under the MacSyFinder v2 framework [21] and HMMER v3.3.2 [26]. Detected systems were further classified by predominant mechanism of action into Abortive infection (Abi), Nucleic acid degradation (NAD; including restriction-modification, CRISPR-Cas, BREX and DdmDE), DNA synthesis inhibition (DSI; including dGTPase, dCTPdeaminase and Viperin) and Other or unknown, following DefenseFinder wiki annotations and the functional literature for each family [20, 27, 28].

### Antimicrobial resistance, virulence factors and prophages

Acquired antimicrobial resistance genes were detected with ABRicate v1.0.1 screened against the NCBI AMRFinder reference database at minimum 90% identity and 80% coverage [29]. The chromosomal intrinsic *V. cholerae* beta-lactamase *varG* was excluded from acquired-AMR counts. These thresholds favour specificity over sensitivity and may under-detect divergent beta-lactamases and novel SXT/R391 variants; the conservative choice was retained to minimise false positives at the cost of a small false-negative penalty. Predicted (genotypic) multidrug resistance was defined as carriage of three or more distinct acquired AMR genes, with the chromosomal intrinsic *varG* excluded. Virulence factors were detected with ABRicate against the VFDB database [30] at the same identity and coverage thresholds, and the canonical seventh-pandemic toxigenic profile was defined as joint presence of *ctxA*, *ctxB* and at least ten genes of the toxin-coregulated pilus operon. Prophages were detected with VirSorter2 v2.2.4 [31] (--include-groups dsDNAphage, ssDNA, --min-score 0.5) and counted at the contig level, and plasmid carriage was inferred from plasmid-associated replicons identified by PlasmidFinder [32] regardless of contig topology. All four content classes were integrated into a single per-isolate matrix used for downstream comparison.

### Machine-learning analyses

To move beyond correlative descriptions and quantify how much information the defence profile carries about clinical multidrug resistance, we applied unsupervised and supervised machine-learning methods to the binary per-isolate defence type matrix (140 systems with prevalence above 1%). Unsupervised structure was recovered with Uniform Manifold Approximation and Projection (UMAP, umap-learn v0.5) [33] in two dimensions with Hamming distance, min_dist = 0.1 and n_neighbors = 30 (random_state = 42), and the resulting embedding was clustered with HDBSCAN [44] using min_cluster_size = 40 and min_samples = 10. Cluster-count stability was assessed across 18 combinations of three seeds, three n_neighbors and two min_cluster_size values, returning between five and nine clusters with a mode of six. Supervised prediction of predicted-MDR status was performed with Random Forest (scikit-learn v1.5, 500 estimators, default max_depth, max_features=’sqrt’, class_weight=’balanced’) and XGBoost (v2.0) classifiers [34, 35].

Three protocols were applied to evaluate generalisability. First, five-fold stratified cross-validation estimated area under the receiver operating characteristic curve (AUC), accuracy, F1 score and average precision. Second, a cross-temporal split trained the classifier on isolates collected up to and including 2005 and tested on isolates from 2006 onwards. Third, a cross-regional split trained on South Asian and African isolates and tested on all other regions. To estimate the minimum number of defence systems needed for accurate classification, we ran recursive feature elimination with cross-validation and an independent grid search over the top-k systems ranked by Random Forest importance. Per-isolate explanations were obtained from a Random Forest fitted to the full dataset using TreeExplainer SHAP values [36], with the positive class corresponding to MDR. All machine-learning analyses were performed in Python v3.10.

### Statistical analyses

Continuous-distribution comparisons used two-sided Mann-Whitney U tests with Cliff’s δ as the effect-size measure, and proportions were compared with two-sided Fisher exact tests. Multiple testing was controlled with the Benjamini-Hochberg false-discovery rate (FDR). Continuous-continuous associations were summarised by Spearman ρ (used for all per-isolate count-vs-count comparisons, including defence-prophage relationships), and co-occurrence between two binary variables was quantified by the Phi coefficient. Temporal trends were estimated within 5-year windows, retaining only windows with at least five isolates per group. To partition the contribution of each mobilome component to defence diversity, we fitted a Poisson generalised linear model with log link in statsmodels, using per-isolate defence subtype count as the outcome and serogroup, year of isolation per decade, isolation source, four region indicator variables (with East Asia as reference), plasmid carriage, prophage count and acquired AMR count as covariates; the model was fitted on the 1,486 isolates with complete metadata, with incidence rate ratios (IRRs) reported with Wald 95% confidence intervals. To identify systems most likely to be physically co-acquired with the SXT/R391 cassette, we constructed a bipartite defence-AMR co-occurrence network restricted to defence types of intermediate prevalence (5 to 95%, n = 49) and AMR genes of intermediate prevalence (2 to 95%, n = 27), and retained edges at BH q < 0.01 and |φ| > 0.4. Co-occurrence interpretations are correlative; physical linkage would require long-read assembly. Statistical comparisons and visualisations were performed in R v4.3 using tidyverse, ggplot2, ggpubr, cowplot, patchwork, VennDiagram and RColorBrewer; per-isolate matrix building, multivariable modelling and the network analysis were performed in Python v3.10 using pandas, NumPy, SciPy, scikit-learn and statsmodels.

## Results

### Pandemic O1 *V. cholerae* carries a broader and mechanistically distinct defence repertoire than the environmental non-O1

We assembled 1,740 *V. cholerae* RefSeq genomes (773 O1, 967 non-O1) sampled between 1934 and 2022 from 60 countries (Fig. S1). The cohort is dominated by cholera-endemic regions, with the largest contributions from Bangladesh, the United States, India and China, and isolates concentrated in clinical sources for O1 and a more balanced clinical-environmental split for non-O1. Geographically, defence diversity was structured with endemic South Asian and African O1 lineages carried the densest defence and acquired AMR repertoires, while the clonal Caribbean Haiti 2010 lineage carried a comparatively depleted repertoire, and prophage carriage in O1 was saturated near 97% across every region (Fig. S2).

Across the 1,740 genomes we detected 396 defence subtypes (141 types, 97 families). O1 carried significantly more subtypes per genome than non-O1 (mean 11.9 vs 9.7; Mann-Whitney P < 0.001, Cliff’s δ = 0.49; Fig. 1a) and more distinct types (median 12 vs 8; δ = 0.64; Fig. 1b), with subtype and type counts tightly coupled in both serogroups (R = 0.95 in O1, R = 0.94 in non-O1; Fig. S3). Despite carrying more systems per genome, O1 contributes only 13 unique subtypes versus 283 unique to non-O1, with 100 subtypes shared (Fig. 1c); 55 entire defence families are absent from any O1 isolate while no family is uniquely O1, establishing the environmental compartment as the principal source of defence diversity in the *V. cholerae* pan-genome.

**Fig. 1.**
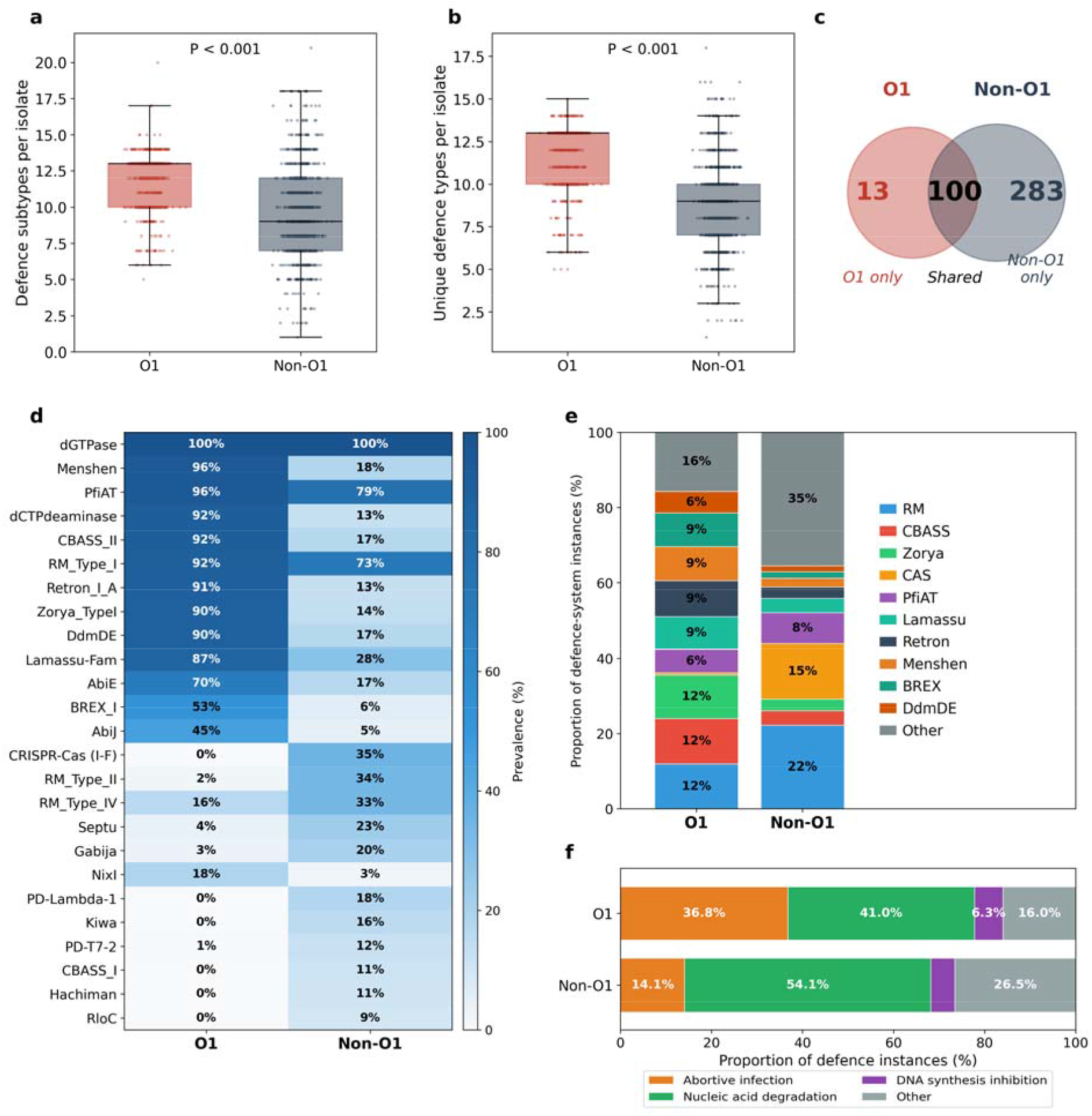
Anti-phage defence diversity in O1 versus non-O1 *V. cholerae*. (a) Per-isolate count of distinct defence subtypes. (b) Per-isolate count of distinct defence types. (c) Heatmap of prevalence of the top 25 most abundant defence types in O1 and non-O1; cells annotated with per-serogroup prevalence. (d) Vertical stacked composition of defence-system instances by family. (e) Venn partition of 396 defence subtypes by serogroup. (f) Mechanism-class composition of defence instances by serogroup.

Of all 141 types, 106 (75.2%) differed between serogroups at FDR < 0.05. DdmDE was present in 90% of O1 versus 17% of non-O1, consistent with its known VPI-2 location [8, 12], and Lamassu-Fam, AbiE, AbiJ, BREX-I, CBASS-II and Menshen were similarly O1-enriched; type II/IV restriction-modification systems, Septu, Gabija and several other families were more prevalent in non-O1 and represent the ancestral environmental repertoire [18]. The per-type prevalence heatmap (Fig. 1d; Fig. S4, Fig. S5) confirms these distinct serogroup signatures. Family-aggregated composition (Fig. 1e) shows that non-O1 concentrates 44% of its instances in a long tail of low-prevalence families, whereas O1 distributes them more evenly across CBASS, Lamassu, Retron, Menshen, Zorya, dCTPdeaminase and DdmDE (each 7 to 8%).

Mechanism-class composition is also qualitatively different (Fig. 1f). O1 carries a more balanced abortive infection and nucleic acid degradation profile (33.5% Abi, 33.3% NAD, 16.4% DSI) while non-O1 is NAD-dominated (43.3%) with much lower Abi representation (14.0%, chi-square = 1,159, P < 0.001), indicating a defence repertoire weighted toward systems that trigger programmed cell death upon infection [27].

Together, pandemic O1 differs from the non-O1 reservoir along every measure examined and greater absolute burden, a restricted unique pool despite that burden, a distinct set of prevalent systems, a more even family composition, and an Abi-biased mechanism profile. This expanded yet narrowly composed defensome raises the question of whether O1 has been subject to greater historical phage predation, or whether the pandemic defence cargo has been co-acquired with other mobile genetic elements; we test the phage-pressure hypothesis directly next.

### Defence content tracks AMR and plasmid burden rather than prophage burden

Prophage carriage was near-universal in O1 (97.4%) and significantly lower in non-O1 (79.8%; OR = 9.51, Fisher P < 0.001; Fig. 2a), and both serogroups carried 0 to 8 prophages per isolate with O1 shifted to slightly higher counts (Fig. 2b). Nevertheless, the per-isolate relationship between defence subtype count and prophage count ran opposite to prediction: within O1 the correlation was significantly negative (Pearson R = −0.15, P < 0.001), while within non-O1 it was weakly positive (R = +0.06, P = 0.07; Fig. 2c). Pooling the two serogroups cancelled these opposing signals (ρ = 0.05), illustrating why stratification is essential. The pattern was robust to isolation source and repeating the analysis separately for clinical (Fig. 2d) and environmental (Fig. 2e) isolates returned the same direction and magnitude.

**Fig. 2.**
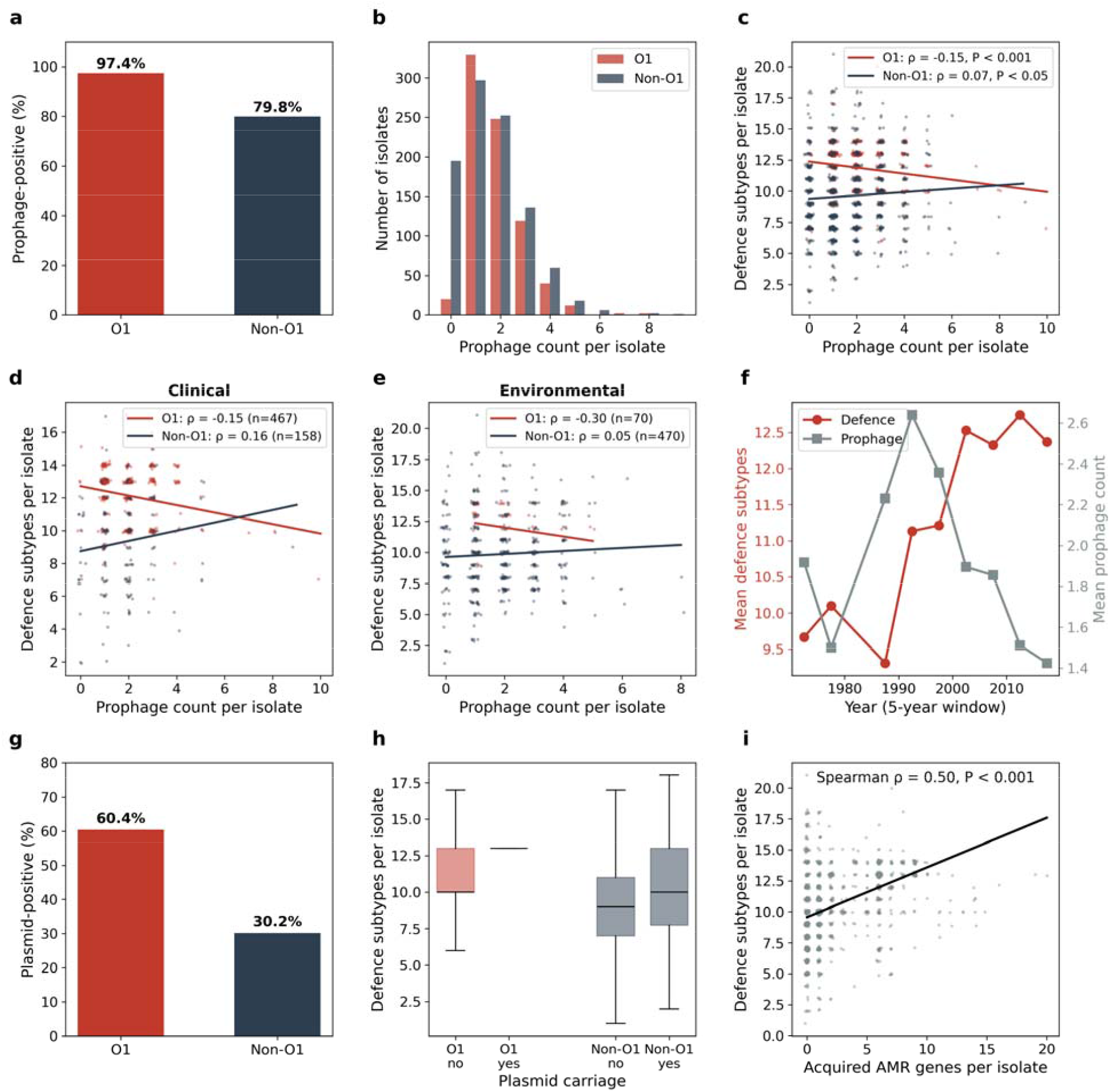
Mobile genetic elements and their relationship to anti-phage defence content. (a) Prophage carriage in O1 and non-O1. (b) Distribution of prophage count per isolate. (c) Defence subtype count versus prophage count in O1 and non-O1. (d) Defence-prophage relationship restricted to clinical isolates. (e) The same analysis restricted to environmental isolates. (f) Mirror temporal trajectories in O1 of mean defence subtype count (red, left axis) and mean prophage count (grey, right axis) per 5-year window from 1970 to 2020. (g) Plasmid carriage in O1 and non-O1. (h) Defence subtype count by plasmid status and serogroup. (i) Defence subtype count versus acquired AMR gene count.

Temporal trajectories provide a second, independent test. Mean defence subtype count in O1 rose from approximately 10 in the 1970s to over 13 by 2010 to 2015, while mean prophage count declined from approximately 2.5 to below 1.5 over the same window (Fig. 2f). These mirror trajectories are the opposite of what sustained phage-predation pressure would predict, ruling out that hypothesis on both contemporary and historical grounds.

Turning instead to the AMR-bearing mobilome, plasmid carriage was twice as high in O1 (60.4%) as in non-O1 (30.2%; OR = 3.53, P < 0.001; Fig. 2g), and plasmid-positive isolates carried more defence subtypes in both serogroups (Fig. 2h). Strikingly, defence subtype count correlated with acquired AMR gene count across the full cohort (Spearman ρ = 0.50, P < 0.001; Fig. 2i) is an order of magnitude stronger than the defence-prophage correlation, implicating the AMR-bearing mobilome as the primary co-traveller of expanded defence content.

### O1 carries a five-fold higher acquired AMR burden dominated by the SXT/R391 cassette

Building on the strong defence-AMR correlation that emerged from the mobilome analysis, we characterised the acquired resistome and virulome of the two compartments in detail to identify which specific resistance genes drive the elevated O1 burden. Median acquired AMR was 5 in O1 versus 1 in non-O1 (Mann-Whitney P < 0.001; Fig. 3a), and 65.2% of O1 met the MDR definition compared with 12.4% of non-O1 (OR = 13.2, P < 0.001; Fig. 3b). The canonical 7PET toxigenic profile, defined as joint carriage of *ctxA*, *ctxB* and complete TCP, was present in 96.9% of O1 and 0% of non-O1, providing near-perfect diagnostic separation between the two compartments (Fig. 3c).

**Fig. 3.**
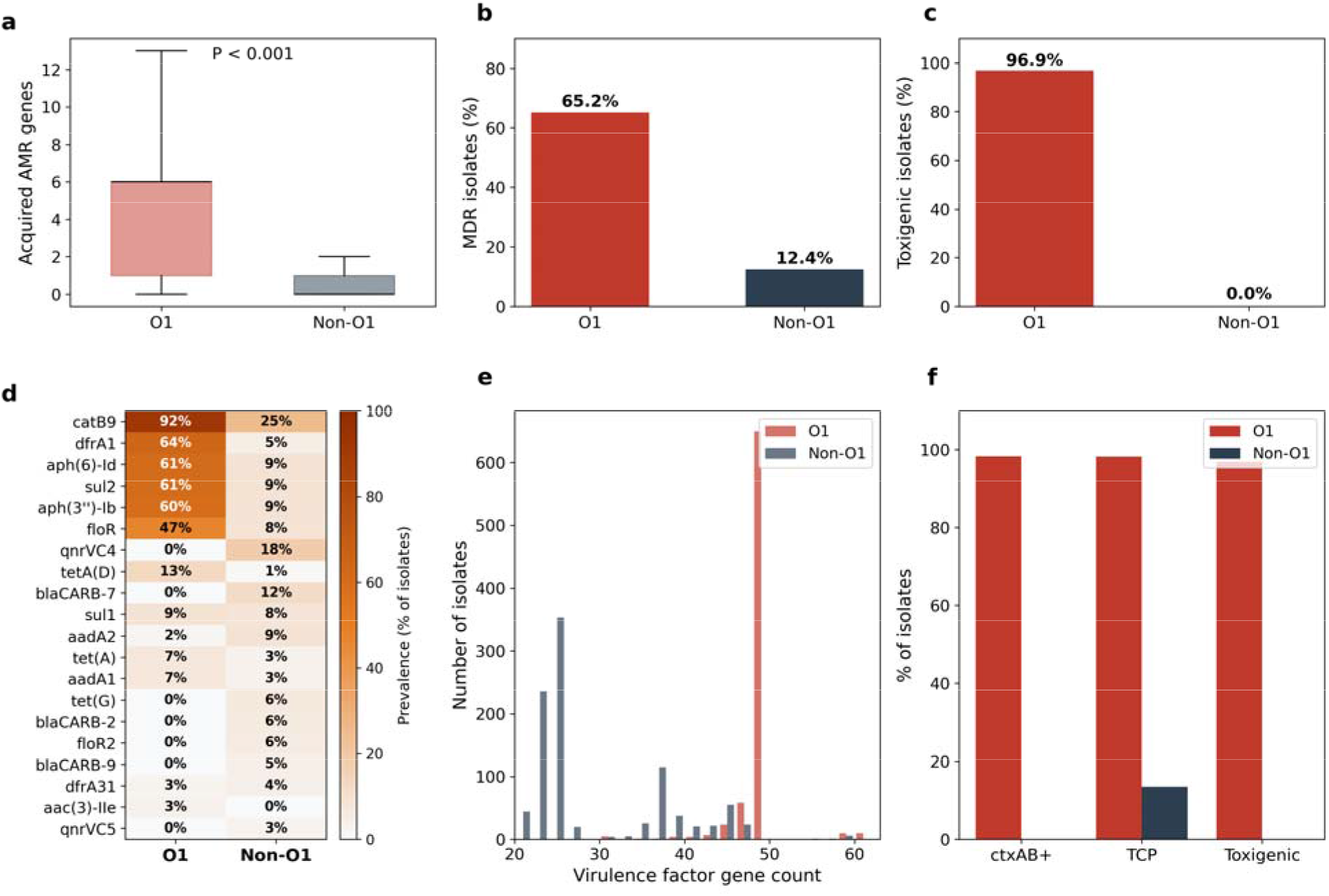
Antimicrobial resistance and virulence-factor landscape of the cohort. (a) Acquired AMR gene count per isolate. (b) Prevalence of multidrug resistance in O1 and non-O1. (c) Prevalence of the canonical seventh-pandemic toxigenic profile. (d) Heatmap of prevalence of the top 20 acquired AMR genes in O1 and non-O1. (e) Virulence-factor gene count distribution. (f) Individual toxigenic markers.

The specific genes responsible for the elevated O1 resistance burden were the canonical SXT/R391 ICE cassette *catB9*, *dfrA1*, *sul2*, *aph(6)-Id*, *aph(3’’)-Ib* and *floR*, each present in 47 to 92% of O1 versus 5 to 25% of non-O1 and each enriched at FDR < 0.001 (Fig. 3d). By contrast, the non-O1 acquired resistome was dominated by *qnrVC4* (18%), *blaCARB-7* (12%) and *tetA*(D) (1%), consistent with environmental acquisition routes distinct from SXT/R391. The virulence-factor compartment showed an equally clear separation: O1 isolates clustered tightly around 48 to 50 VF genes, while non-O1 spanned a much broader distribution from 20 to 60 genes (Fig. 3e), and the individual toxigenic markers *ctxAB*+, complete TCP and the composite toxigenic profile each separated the two serogroups close to perfectly (Fig. 3f).

Overall, the AMR and virulence-factor landscapes establish that pandemic O1 carries a quantitatively and qualitatively distinct mobilome cargo dominated by the SXT/R391 ICE cassette. The strong correlation between defence and AMR content from the preceding section, combined with the present finding that O1 AMR is carried predominantly on a single well-characterised ICE family, predicts that defence systems could display specific co-occurrence patterns with individual SXT/R391 cassette genes. We test this prediction explicitly in the network analysis in following section but first place the AMR landscape to determine the temporal pattern.

### Defence and prophage burden follow mirror trajectories with the SXT/R391 acquisition window

Having established the temporal defence-AMR association, we asked when this association was established across the eight-decade span of the cohort. Median defence subtype count per 5-year window rose in O1 from 10 in the 1970s to 13 by 2010 to 2015 (R = 0.45, P < 0.001), while non-O1 fluctuated around 9 throughout (Fig. 4a). Median acquired AMR per window in O1 increased sharply between 1995 and 2000 from a baseline of 1 to a plateau of 6 (Fig. 4b). Plasmid carriage in O1 rose from below 25% before 1990 to over 80% by 2000 (Fig. 4c), and prophage carriage in O1 remained near saturation throughout while non-O1 rose gradually over the same period (Fig. 4d). These coordinated trajectories indicate that the major shifts in the O1 mobilome compartment occurred during a narrow window between approximately 1990 and 2005.

**Fig. 4.**
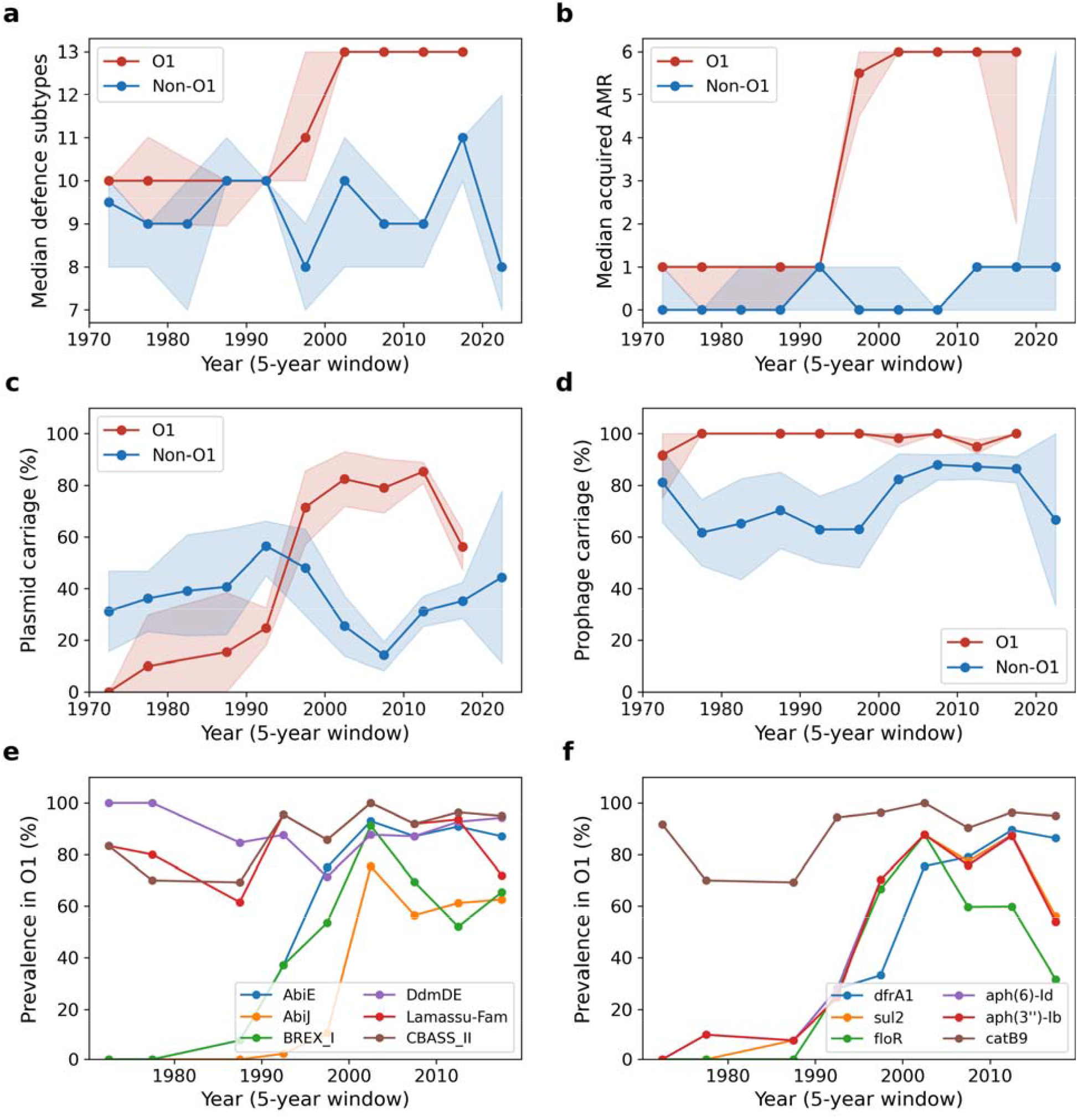
Temporal dynamics of defence and resistance content in *V. cholerae*. The data is from 1970 to 2022 and summarised in 5-year windows. Shaded bands in panels (a) to (d) show 95% bootstrap confidence intervals (200 resamples). (a) Median defence subtype count per window. (b) Median acquired AMR per window. (c) Plasmid carriage in O1 and non-O1. (d) Prophage carriage in O1 and non-O1. (e) Prevalence of six selected defence systems in O1. (f) Prevalence of the six canonical SXT/R391 ICE resistance genes in O1.

Examining individual systems within this window confirmed the coordinated nature of the shift. AbiJ rose from undetected before 1985 to 76% by the early 2000s, and BREX-I from below 10% to approximately 95% over the same window (Fig. 4e), consistent with the SXT-ICE BREX hotspot reported by LeGault et al. [18] and Adams et al. [12]. Family- and mechanism-level temporal trajectories exhibit similar patterns at a broader resolution (Fig. S6, Fig. S7). The canonical SXT/R391 ICE resistance genes *catB9*, *dfrA1*, *sul2*, *aph(6)-Id*, *aph(3’’)-Ib* and *floR* rose in parallel during 1995 to 2000 (Fig. 4f), placing the AMR acquisition window in lockstep with the defence acquisition window.

Our 1934 to 2022 reconstruction demonstrates that defence and prophage have followed mirror trajectories in pandemic *V. cholerae* for at least four decades, with the rise of defence systems temporally locked to the global emergence of SXT/R391 ICE-borne AMR. The temporal coincidence of defence and AMR expansion is consistent with shared horizontal-transfer routes but does not by itself identify which specific defence systems are most likely to be co-acquired with which AMR genes. We address that question with a population-scale defence-AMR co-occurrence network in the next section.

### Network and multivariable analyses nominate specific defence systems as candidate co-occurring partners of the SXT/R391 cassette

The temporal coincidence between defence and AMR expansion suggests direct genetic linkage. To test this at genome-pair resolution we constructed a population-scale defence-AMR co-occurrence network: Phi correlations between defence types of intermediate prevalence (5 to 95%, n = 49) and AMR genes of intermediate prevalence (2 to 95%, n = 27) were computed and filtered at FDR-adjusted |φ| > 0.4, retaining 56 significant positive pairs (Fig. 5a-b). The strongest associations were AbiE-*dfrA1* (φ = 0.80), dCTPdeaminase-*catB9* (φ = 0.75), AbiE-*aph(6)-Id* (φ = 0.74), BREX-I-*floR* (φ = 0.73) and AbiJ-*dfrA1* (φ = 0.73). These associations held under joint adjustment for serogroup, decade, region, isolation source, plasmid carriage and prophage count in a multivariable Poisson model. O1 serogroup (IRR = 1.17, 95% CI 1.12 to 1.22, P < 0.001), decade (IRR = 1.03, P < 0.001), plasmid carriage (IRR = 1.06, P < 0.01) and AMR gene count (IRR = 1.02, P < 0.001) each remained independent predictors while prophage count did not (P = 0.77).

**Fig. 5.**
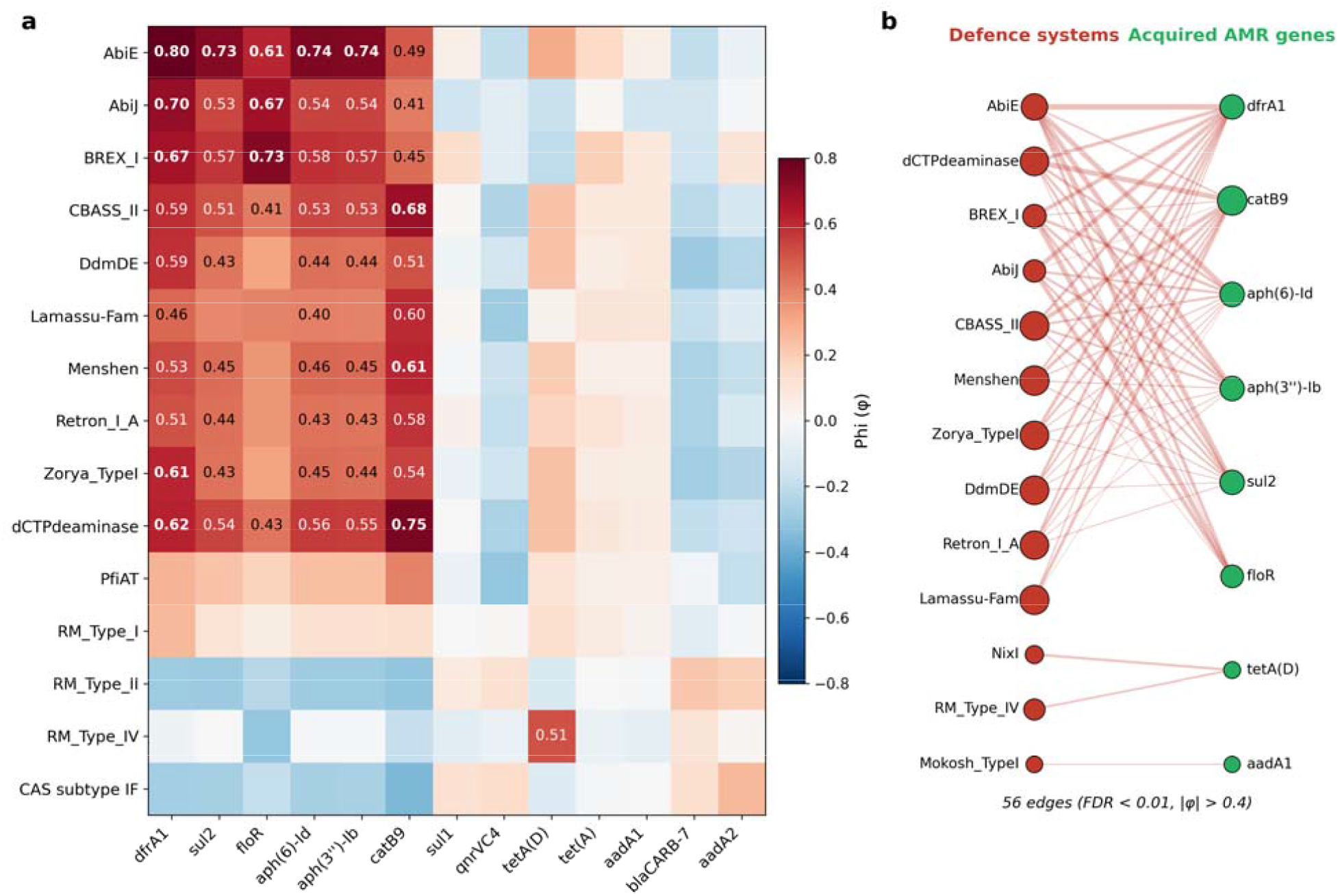
Population-scale defence-AMR co-occurrence structure. (a) Phi coefficient heatmap of 15 defence types versus 13 AMR genes, with cells annotated where |φ| ≥ 0.4. (b) Bipartite network of 56 defence-AMR pairs surviving the FDR q < 0.01 and |φ| > 0.4 thresholds; defence systems are shown as red nodes on the left and acquired AMR genes as green nodes on the right, with node size proportional to within-cohort prevalence and edge thickness proportional to the Phi value.

Several lines of evidence reinforce the network. PCA of defence-type repertoires cleanly separates O1 from non-O1 along PC1 with secondary structure by year and region (Fig. S8). Mosaic plots of the four highest Phi pairs visualise per-isolate co-occurrence patterns (Fig. S9), and within-O1 co-occurrence analysis identifies two anticorrelated defence modules (Fig. S10), a pandemic core (RM-Type I, Zorya-TypeI, DdmDE, AbiE, AbiJ, BREX-I, Lamassu-Fam, dCTPdeaminase, CBASS-II, Retron-I_A, Menshen) and an alternative module (RM-Type II/IV, Septu, Druantia-III, Gabija, NixI, Mokosh-TypeI, PrrC), consistent with distinct integrative-element lineages each carrying a characteristic cassette. The BREX-I-floR association recapitulates the mechanistically validated SXT-ICE BREX hotspot [18], providing internal validation; the remaining strong associations involve systems (AbiE, AbiJ, dCTPdeaminase, Menshen, Retron-I_A, Lamassu-Fam) not previously linked to the SXT/R391 ICE, whose effect sizes approach the binary-variable maximum and are more consistent with physical co-linkage than with strain-level co-selection.

### Machine learning quantifies the predictive value of the defensome

Unsupervised UMAP projection of the 140-system binary defence matrix resolved a clear O1 versus non-O1 separation, and HDBSCAN clustering identified seven distinct defensome lineages (Fig. 6a-b). UMAP and HDBSCAN stability across seeds and parameter settings is shown in Fig. S11. The largest cluster (cluster 4; n = 878, 3.4% O1) captured the environmental non-O1 compartment with a median AMR burden of 1.2 and MDR rate of 12%. The four O1-dominated clusters (91 to 100% O1) carried mean AMR burdens of 5.7 to 7.0 and MDR rates of 83 to 98%, with early (cluster 0, median year 1994) and modern (cluster 3, median year 2013) 7PET sub-lineages provisionally distinguishable by sampling year. A small cluster (n = 180, 85% O1, median year 1994, MDR rate 11%) captured early O1 isolates that had not yet acquired the SXT/R391 defence cassette, structurally confirming the 1990 to 2000 acquisition window from the temporal analysis; MDR rate scaled near-linearly with mean AMR burden across clusters (Fig. 6c).

**Fig. 6.**
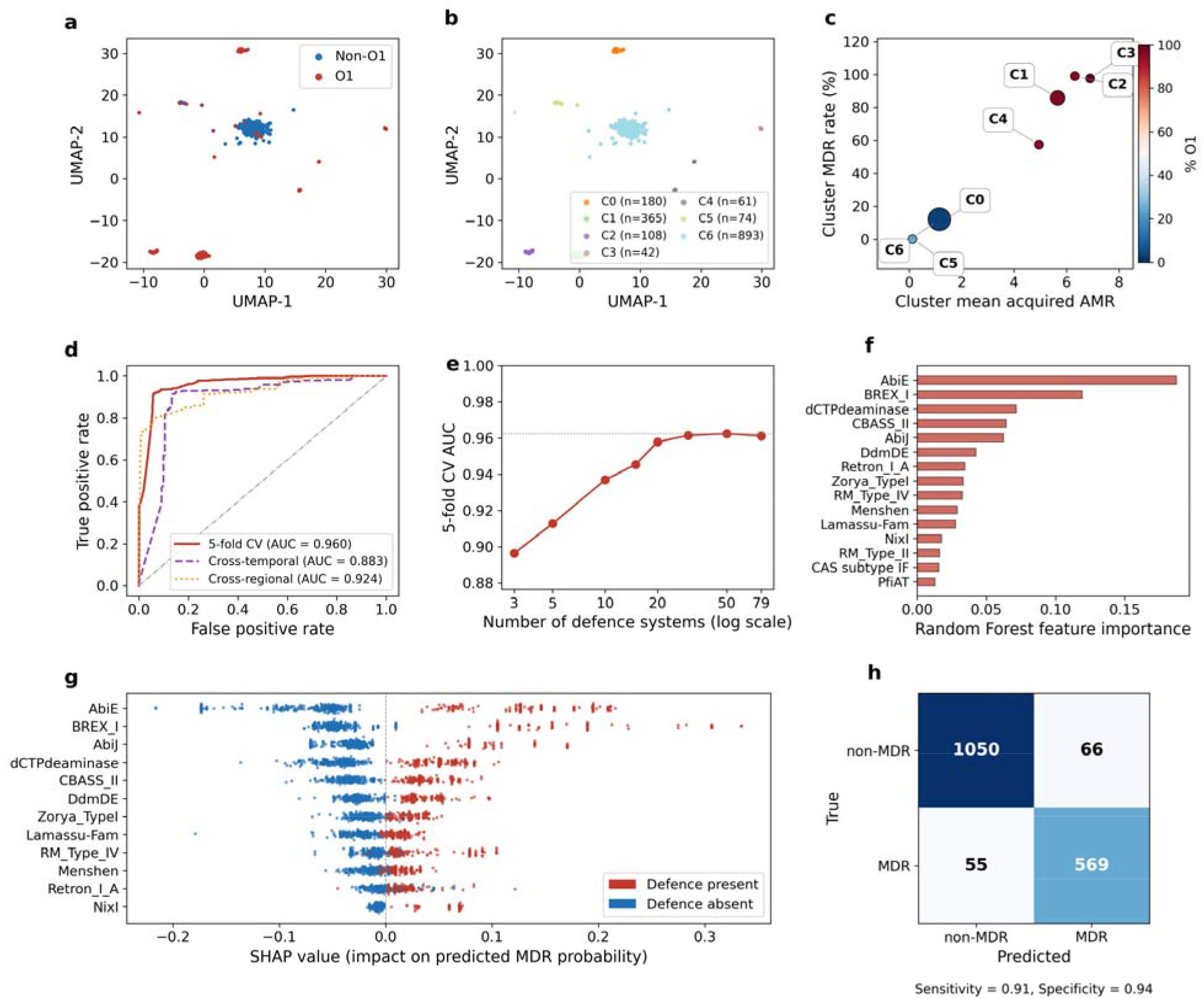
Machine-learning analyses of the *V. cholerae* defensome. (a) UMAP projection of the 140-system defence matrix coloured by serogroup. (b) UMAP coloured by HDBSCAN cluster. (b) Cluster mean acquired AMR versus cluster MDR rate; point size scales with cluster size and colour with % O1. (d) Receiver operating characteristic curves under three protocols, 5-fold cross-validation, cross-temporal, and cross-regional (e) 5-fold CV AUC as a function of the number of defence systems used (f) Top 15 Random Forest feature importances. (g) SHAP summary plot for the top 12 defence systems. (h) Confusion matrix at the 0.5 probability threshold.

A Random Forest classifier trained on the defence profile alone achieved a pooled 5-fold cross-validated AUC of 0.96 (Fig. 6d). Performance generalised across time (out-of-time AUC 0.88, AUPR 0.78; trained pre-2005, tested post-2005) and geography (cross-regional AUC 0.92, AUPR 0.86; trained South Asia and Africa, tested elsewhere). Within-serogroup refitting showed the signal exceeds the O1/non-O1 composition difference, within-O1 AUC 0.92, within-non-O1 AUC 0.93, versus a serogroup-only logistic baseline of 0.78. Ablating the six most O1-enriched defence types results in pooled AUC at 0.96 (Fig. S12), and recursive feature elimination showed that five systems sufficed for AUC = 0.91 while ten reached AUC = 0.94 (Fig. 6e). The top-ranked features by Random Forest importance (AbiE, BREX-I, CBASS-II, dCTPdeaminase, AbiJ; Fig. 6f) recapitulated the network Phi ranking almost exactly, and SHAP per-isolate explanations confirmed that presence of each top-12 system consistently pushed predicted MDR probability upwards (Fig. 6g). At the standard 0.5 threshold the classifier achieved 91.0% sensitivity and 94.3% specificity (Fig. 6h).

Together, these results show that the defensome carries near-diagnostic information about MDR status, that this predictive structure generalises across decades and geography, and that a compact five-system signature (AbiE, BREX-I, CBASS-II, dCTPdeaminase, AbiJ) recovers most of the available signal, motivating prospective evaluation of the defensome as a low-cost genomic correlate of multidrug resistance.

## Discussion

Our pan-genomic analysis of 1,740 *V. cholerae* genomes sampled between 1934 and 2022 addresses three gaps in the current understanding of the *V. cholerae* defensome and places its findings in the context of the broader SXT/R391 ICE biology. First, the environmental non-O1 compartment harbours 283 unique defence subtypes and 55 unique families that are entirely absent from pandemic O1 genomes, establishing the environmental reservoir as the dominant source of anti-phage immune diversity in the *V. cholerae* pan-genome. This scale of non-O1-unique diversity is consistent with the emerging view that chromosomal integrons and horizontally mobilised elements serve as biobanks from which bacterial lineages can acquire novel defence cassettes [14], and with earlier proposals that non-pandemic *V. cholerae* may harbour defence strategies not yet encountered in pandemic lineages [19]. Despite carrying more defence systems per genome on average, O1 contributes only 13 unique subtypes against the 283 contributed by non-O1, a disproportion that underscores how the pandemic clade has expanded its defence content by repeatedly drawing from a shared or non-O1-derived pool rather than by diversifying independently.

Second, the relationship between defence diversity and prophage burden is fundamentally different between the two compartments and is incompatible with a simple phage-predation selectionist model. Within O1 the correlation is significantly negative, within non-O1 it is weakly positive, and the pooled estimate is negligible, a pattern that illustrates Simpson’s paradox and would have been missed without serogroup stratification. The temporal reconstruction extends this argument across five decades: rising defence diversity in O1 is mirrored by declining rather than rising prophage burden, and prophage count retains no independent explanatory power once serogroup, decade and mobilome covariates are included in the multivariable model. These converging lines of evidence indicate that horizontal transfer via the AMR-bearing mobilome, rather than selection driven by phage predation, is the primary force shaping O1 defence content [37, 38].

Third, the 88-year temporal reconstruction shows that the rise of specific defence systems in O1 was temporally locked to the emergence of SXT/R391 ICE-borne resistance genes during a narrow window between approximately 1990 and 2005. The prevalence of *abiJ* and *BREX-I* rose from below 10% before 1985 to above 75% by the early 2000s, closely paralleling the acquisition of *catB9*, *dfrA1*, *sul2* and *floR* on the canonical SXT/R391 cassette. This temporal coincidence provides population-scale, long-resolution support for the defence-AMR co-transfer principle demonstrated through clinical sampling [18] and reviewed in [16, 17]. The bipartite network analysis shows that the co-dissemination signature extends beyond the established BREX and restriction-modification hotspot. Six systems, AbiE, AbiJ, dCTPdeaminase, Menshen, Retron-I_A and Lamassu-Fam, each display Phi associations above 0.65 with multiple SXT/R391 cassette AMR genes, with AbiE-*dfrA1* reaching φ = 0.80. The strong co-occurrence of DdmDE and dCTPdeaminase with SXT/R391 AMR genes most likely reflects linkage disequilibrium within the phylogenetically constrained 7PET background, given their established positions on VPI-2 and a VSP-II variant [10, 39]. The remaining candidates, AbiE, AbiJ, Retron-I_A, Lamassu-Fam and Menshen, have not previously been linked to SXT/R391 or the core pandemic islands and are high-priority targets for long-read assembly and functional characterisation.

The machine-learning analyses translate the correlative network findings into a quantitative predictive framework. A Random Forest classifier trained on the 140-system binary defence profile achieves a pooled cross-validated AUC of 0.96, and performance is preserved across time (cross-temporal AUC 0.88) and geography (cross-regional AUC 0.92), confirming that the predictive signal is biological rather than an artefact of training-set composition. Recursive feature elimination identifies a compact five-system signature, AbiE, BREX-I, CBASS-II, dCTPdeaminase and AbiJ, sufficient for AUC of 0.91. The seven UMAP-defined defensome lineages capture both the pre-acquisition phase (cluster 6, median year 1994, MDR rate 11%) and the post-acquisition phase (clusters 0 to 3, MDR rates 83 to 98%), providing structural support for the co-transfer interpretation. These findings suggest that a small panel of DefenseFinder calls could serve as a low-cost genomic flag for multidrug resistance in surveillance settings where targeted PCR is unavailable [40, 41], though prospective validation in phenotyped cohorts is needed before clinical application.

The mechanism-class analysis adds a further dimension. O1 carries a more balanced abortive infection and nucleic acid degradation repertoire than non-O1 (33.5% Abi versus 14.0% Abi), which may reflect selection for cell-altruistic immunity strategies within the clonal 7PET expansion [2]. This compositional shift has practical implications for phage therapy directed at circulating O1 lineages, as Abi-dominated defensomes may limit the efficacy of single-phage approaches and favour receptor-diverse multiphage formulations [42].

In summary, this study provides the largest automated *V. cholerae* defensome resource to date, identifies the environmental non-O1 reservoir as a far richer source of defence diversity than previously appreciated, and delivers population-scale temporal evidence that pandemic defence content has been co-acquired with AMR as cargo on SXT/R391-family ICEs. The defensome carries near-diagnostic information about multidrug resistance that generalises across decades and geography, supporting its prospective evaluation as a surveillance tool pending phenotypic validation.

## Author contributions

MMH, DM, JSM and GEV designed the study. MMH, JT, DL and GEV performed the analyses and interpreted the data. MMH took the lead to write the manuscript, bioinformatic and machine learning analyses. DM and GEV provided funding. All authors reviewed and provided critical feedback of the manuscript.

## Conflicts of interest

The authors declare that there are no conflicts of interest.

## Funding information

This study was supported by Australian Research Council Discovery Projects DP230101760 and DE250100444, the Australian Institute for Microbiology and Infection (AIMI), Faculty of Science, University of Technology Sydney.

## Notes

### Competing Interest Statement

The authors have declared no competing interest.

