## Supplemental File for "Co-dissemination of anti-phage defence systems and multidrug resistance is associated with the seventh-pandemic *Vibrio cholerae* defensome"

### Contents

This supplementary file contains 12 supplementary figures (S1 to S12).

### Supplementary Figures

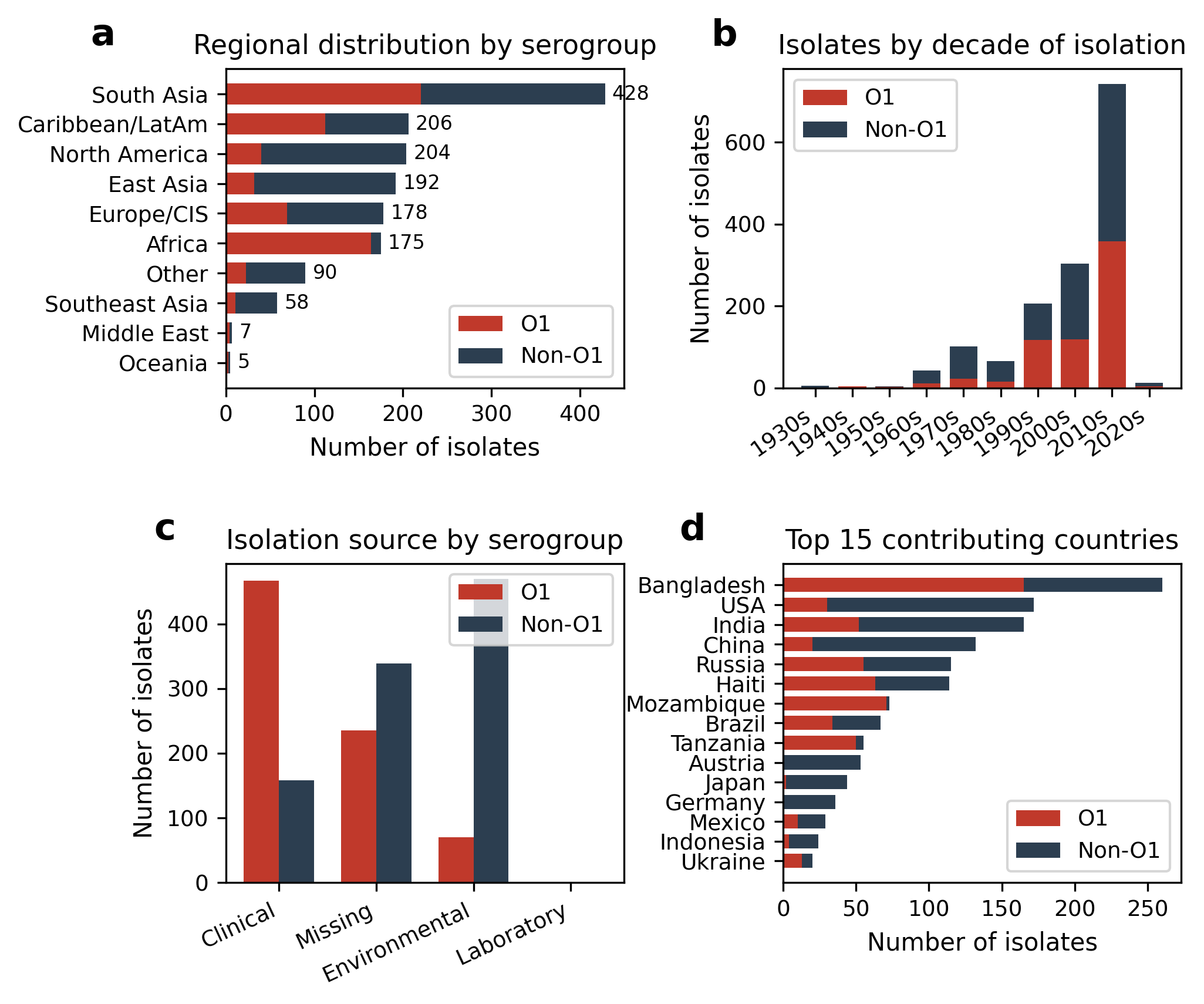

**Fig. S1. Cohort overview of the 1,740 *V. cholerae* genome collection.** (a) Regional distribution of isolates by serogroup. (b) Distribution of isolates by decade of isolation. (c) Distribution by isolation source by serogroup. (d) Top 15 contributing countries by serogroup.

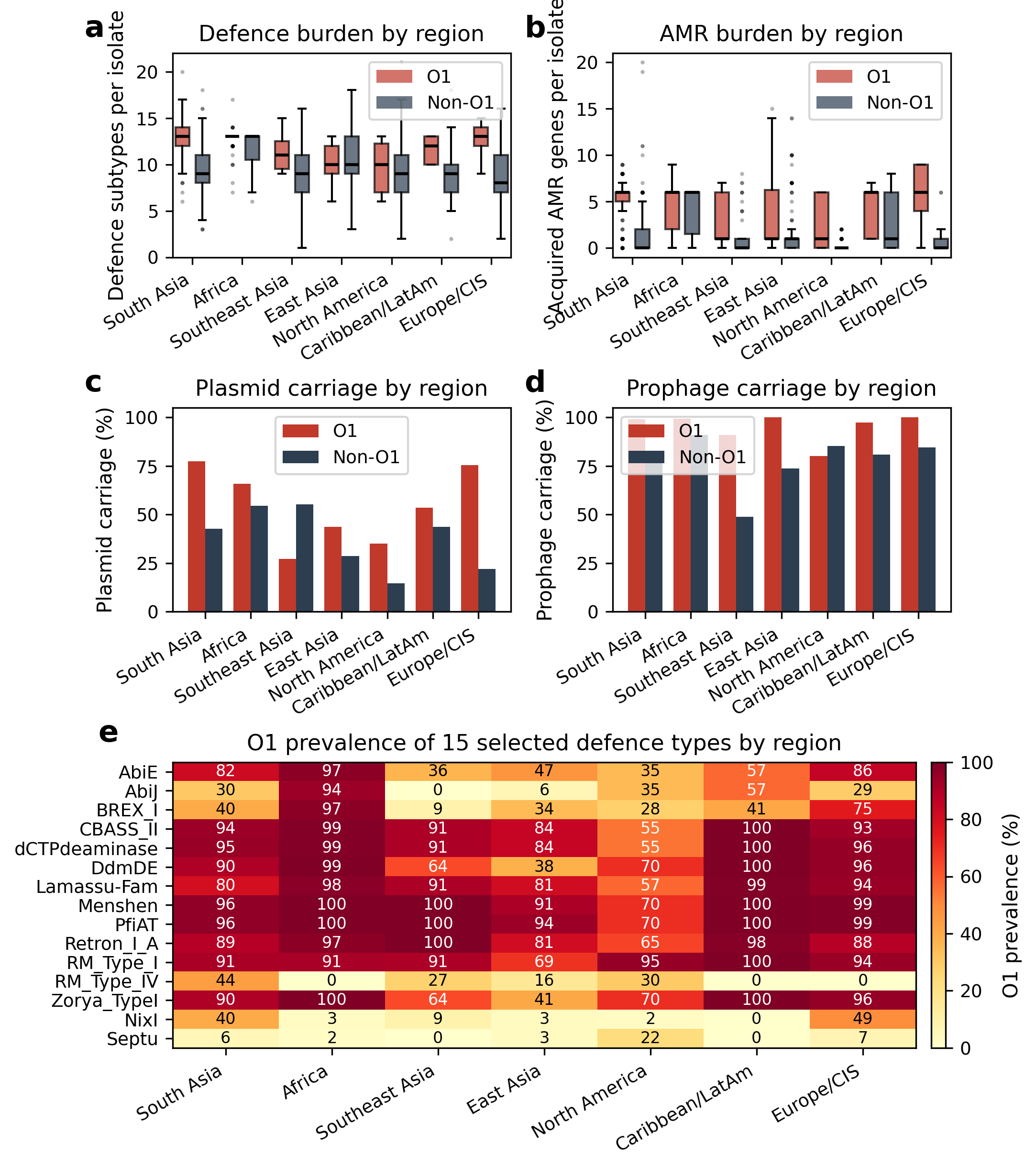

**Fig. S2.** **Spatial distribution of defence systems and mobile genetic elements across the seven major sampling regions.** (a) Defence subtypes per isolate by region and serogroup. (b) Acquired AMR by region and serogroup. (c, d) Plasmid and prophage carriage by region. (e) Heatmap of O1 prevalence of 15 selected defence types across seven regions.

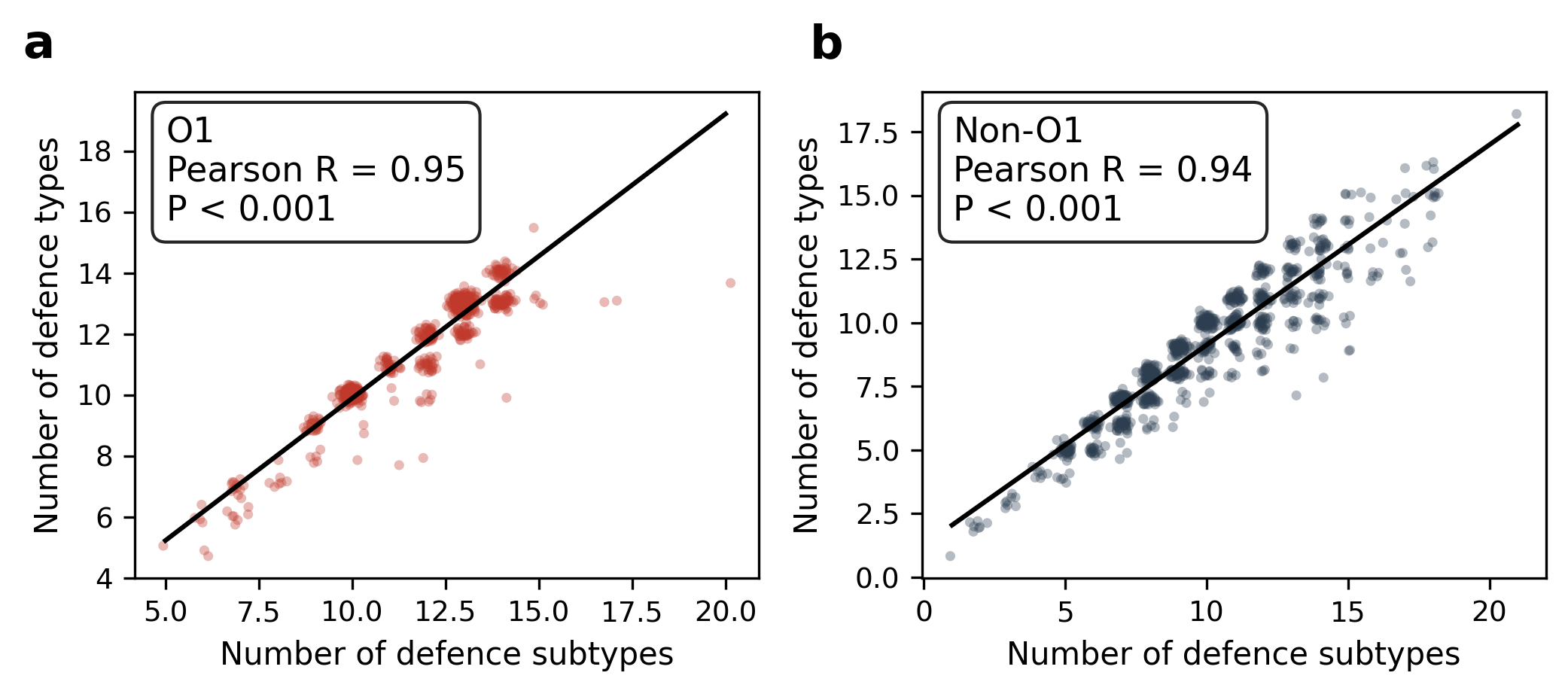

**Fig. S3. Correlation between defence subtype and type counts per isolate, stratified by serogroup.** (a) O1 (Pearson R = 0.95). (b) Non-O1 (Pearson R = 0.94).

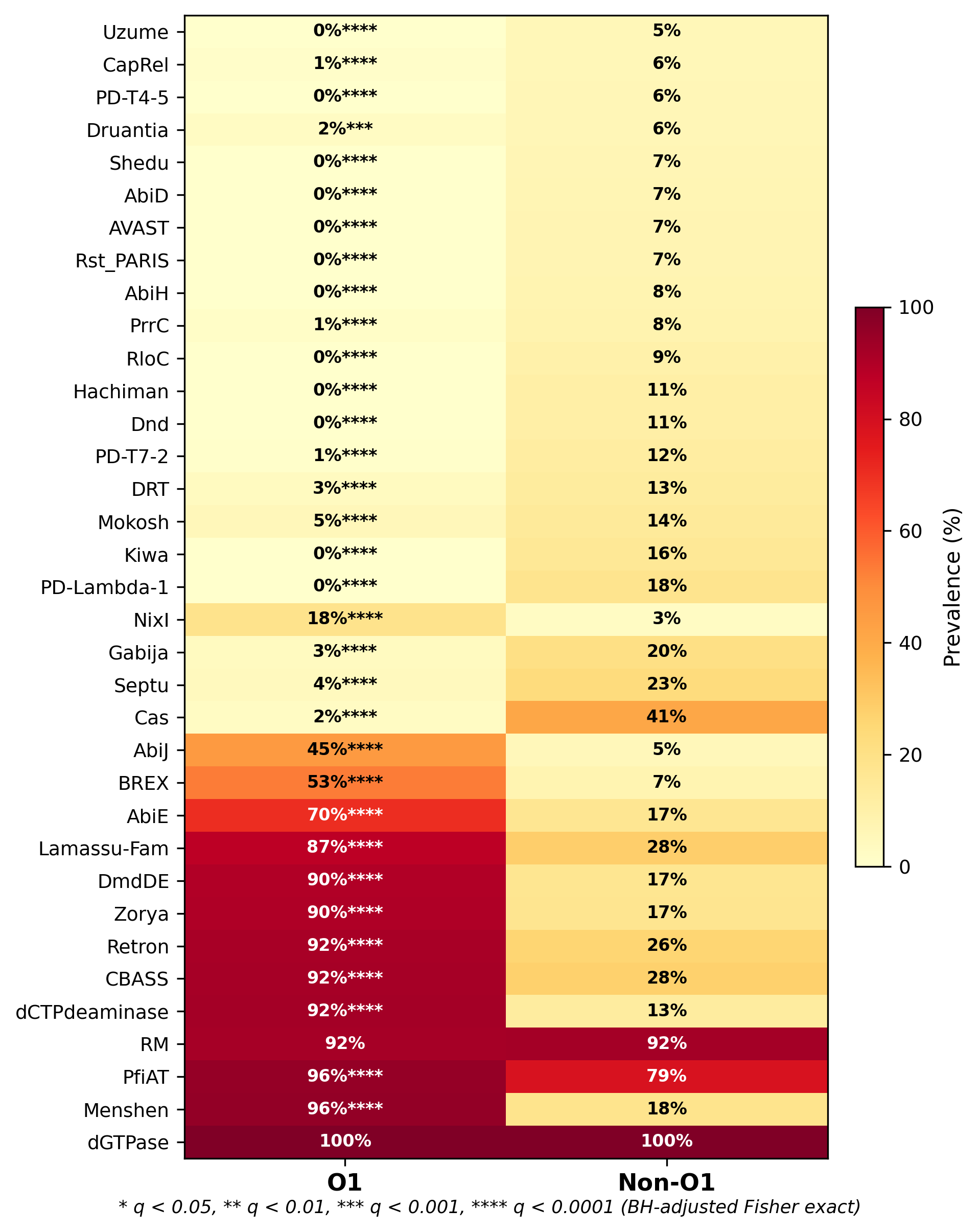

**Fig. S4.** **Heatmap of family-level defence-system prevalence in O1 versus non-O1, showing the top 35 families by overall cohort prevalence.** Cells are annotated with per-serogroup prevalence (%); asterisks denote BH-adjusted q values (* q < 0.05; ** q < 0.01; *** q < 0.001; **** q < 0.0001). The figure complements the type-level view in main Fig. 1c.

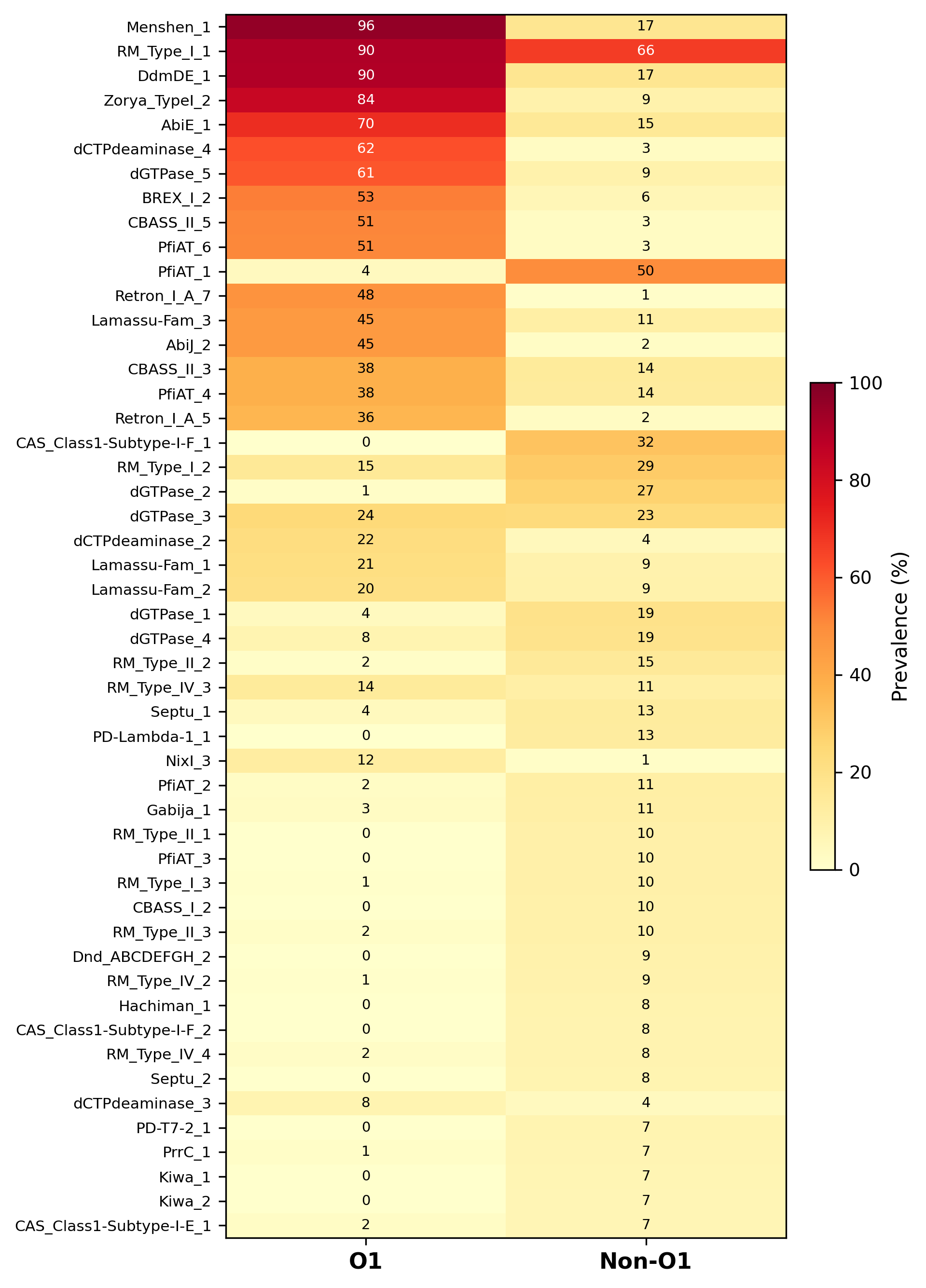

**Fig. S5.** **Compact heatmap of the top 50 defence subtypes by maximum prevalence across O1 and non-O1.** Subtypes are ordered by maximum prevalence, and cells are annotated with per-serogroup prevalence (%).

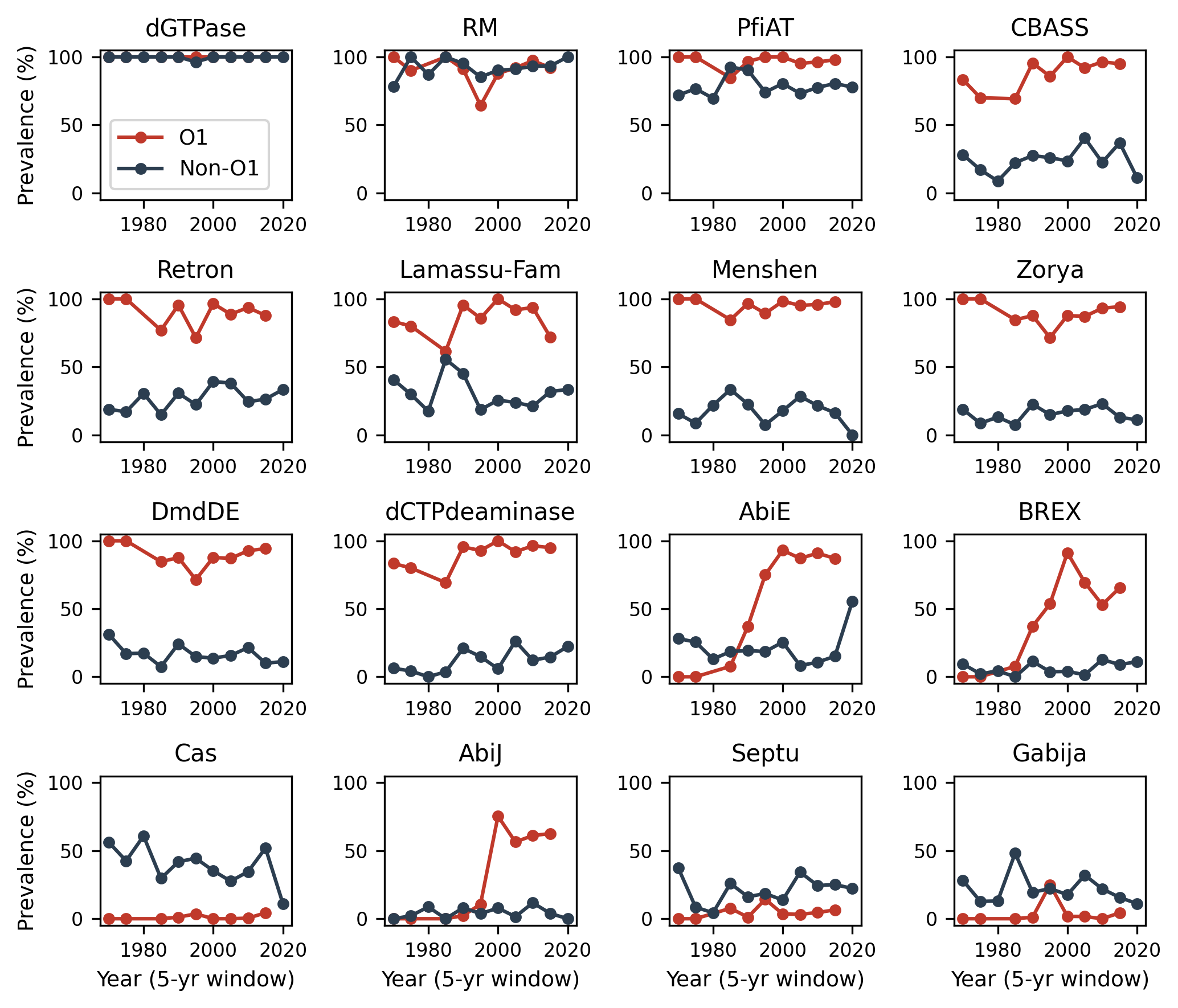

**Fig. S6.** **Temporal trajectories of the most prevalent defence families across 1970 to 2020 in 5-year windows, with O1 (red) and non-O1 (blue) shown per panel.**

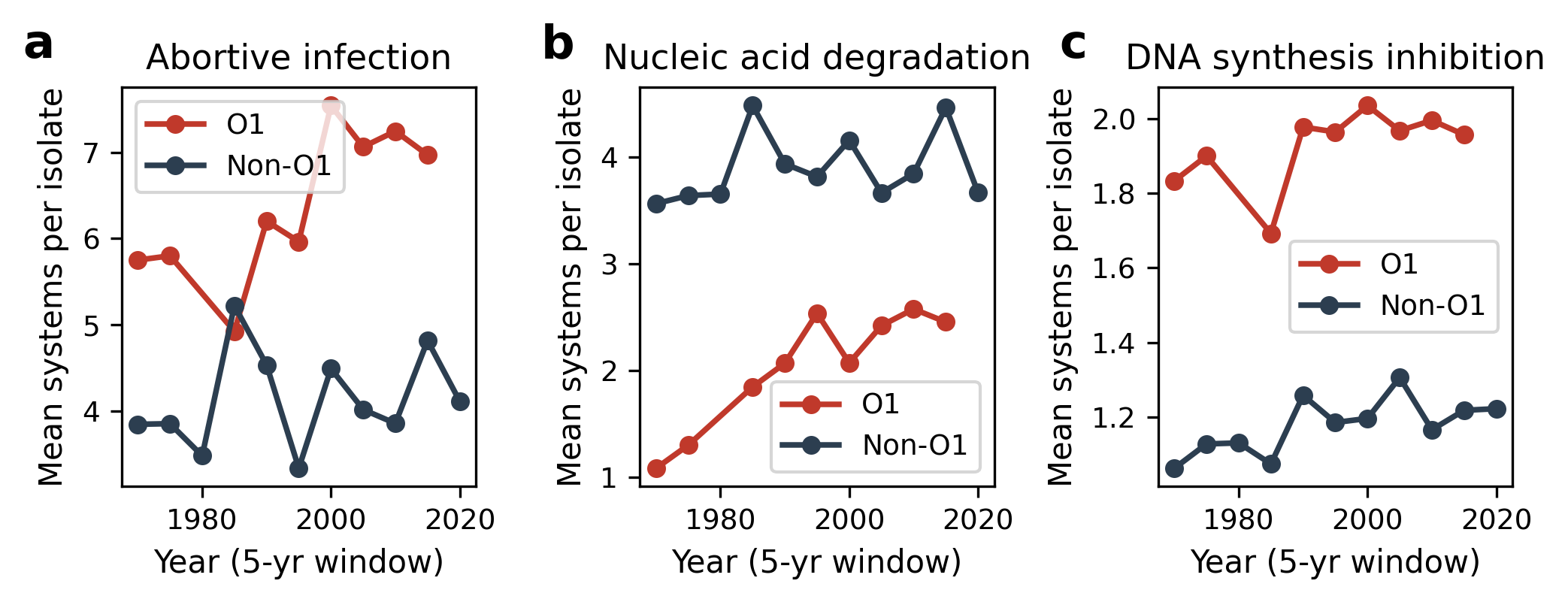

**Fig. S7.** **Per-mechanism temporal trends across 1970 to 2020.** (a) Abortive infection. (b) Nucleic acid degradation. (c) DNA synthesis inhibition.

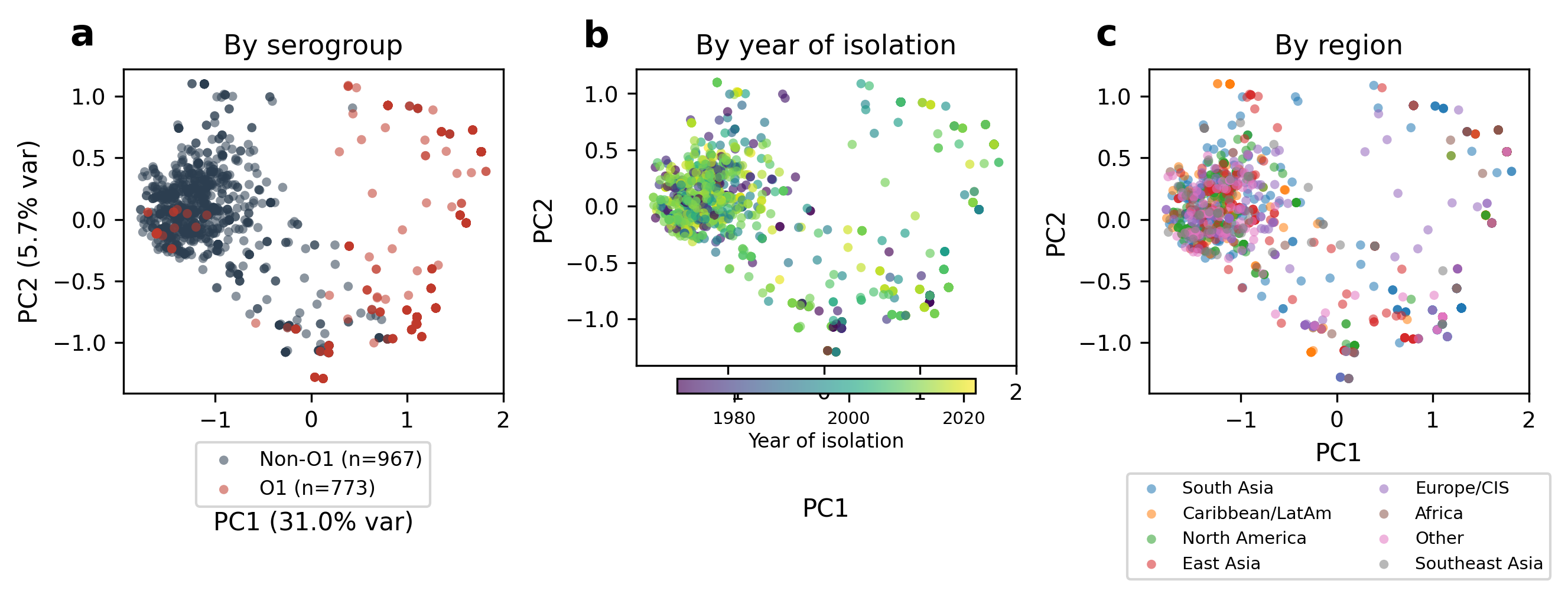

**Fig. S8.** **Principal component analysis of defence-type repertoires.** (a) Coloured by serogroup. (b) Coloured by year of isolation. (c) Coloured by region.

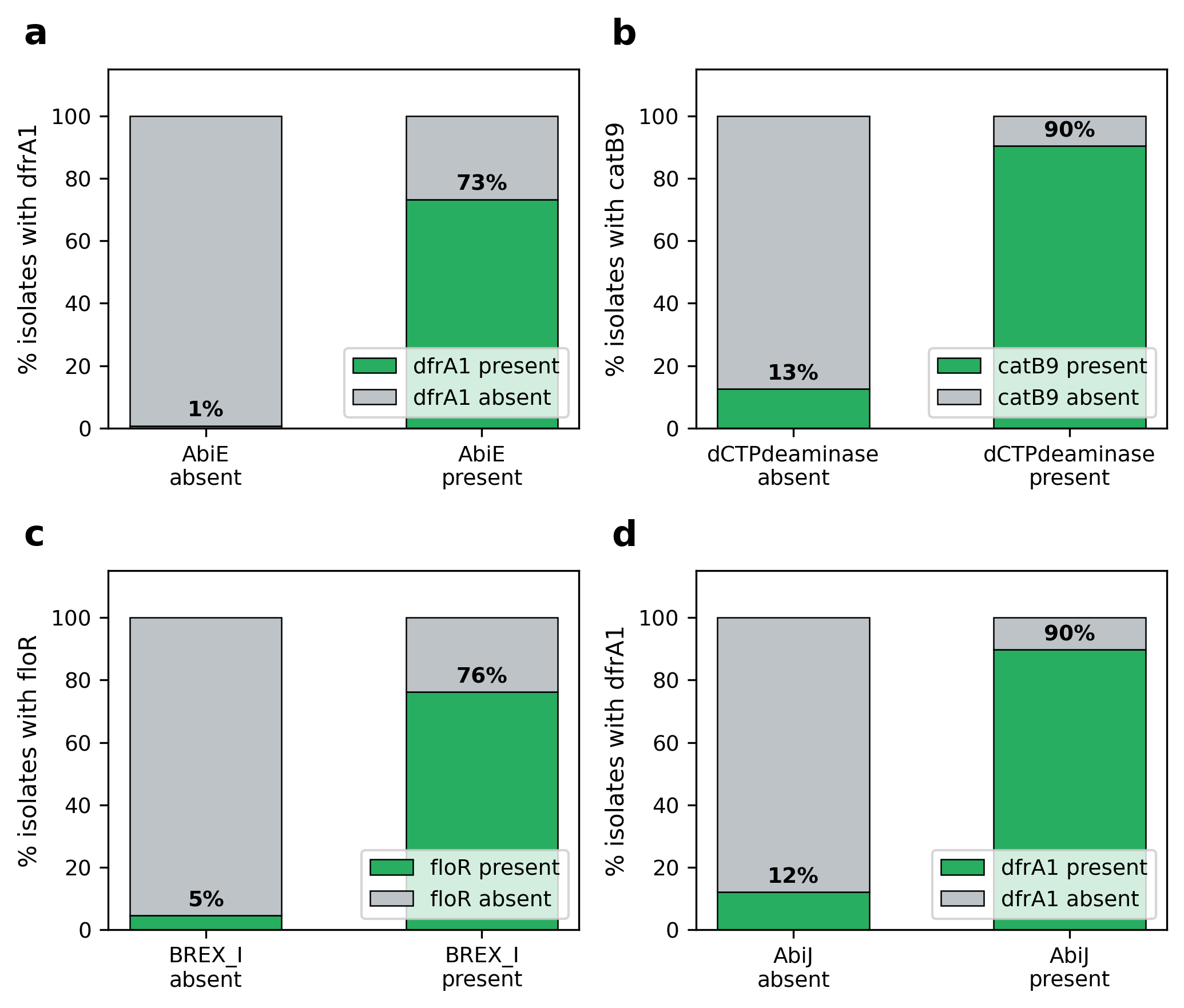

**Fig. S9. Highlighted defence-AMR co-occurrence pairs from the bipartite network analysis.** Each panel shows the percentage of isolates carrying the AMR gene when the defence system is absent (left bar) versus present (right bar). (a) AbiE x dfrA1 (phi = 0.80, P < 0.001). (b) dCTPdeaminase x catB9 (phi = 0.75, P < 0.001). (c) BREX-I x floR (phi = 0.73, P < 0.001). (d) AbiJ x dfrA1 (phi = 0.73, P < 0.001). The figure shows that co-occurrence of each defence system with its partner AMR gene approaches saturation (73 to 90% of defence-present isolates carry the AMR gene), while absence of the defence system is associated with near-absence of the resistance gene.

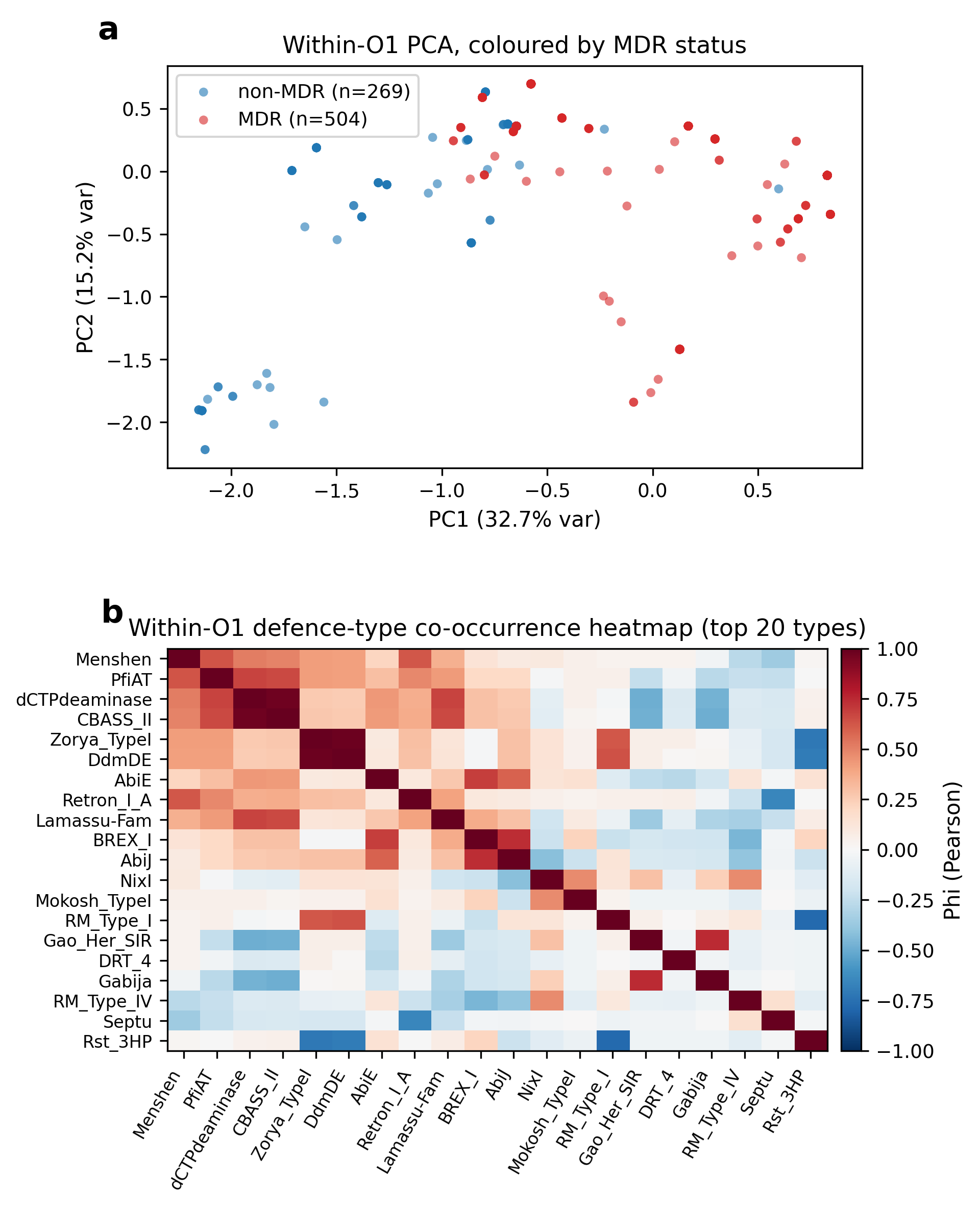

**Fig. S10. Joint co-evolution of defence and resistance across the cohort.** (a) Within-O1 PCA on the defence-type matrix coloured by MDR status. (b) Within-O1 defence-type co-occurrence heatmap identifying a pandemic core module and an alternative module.

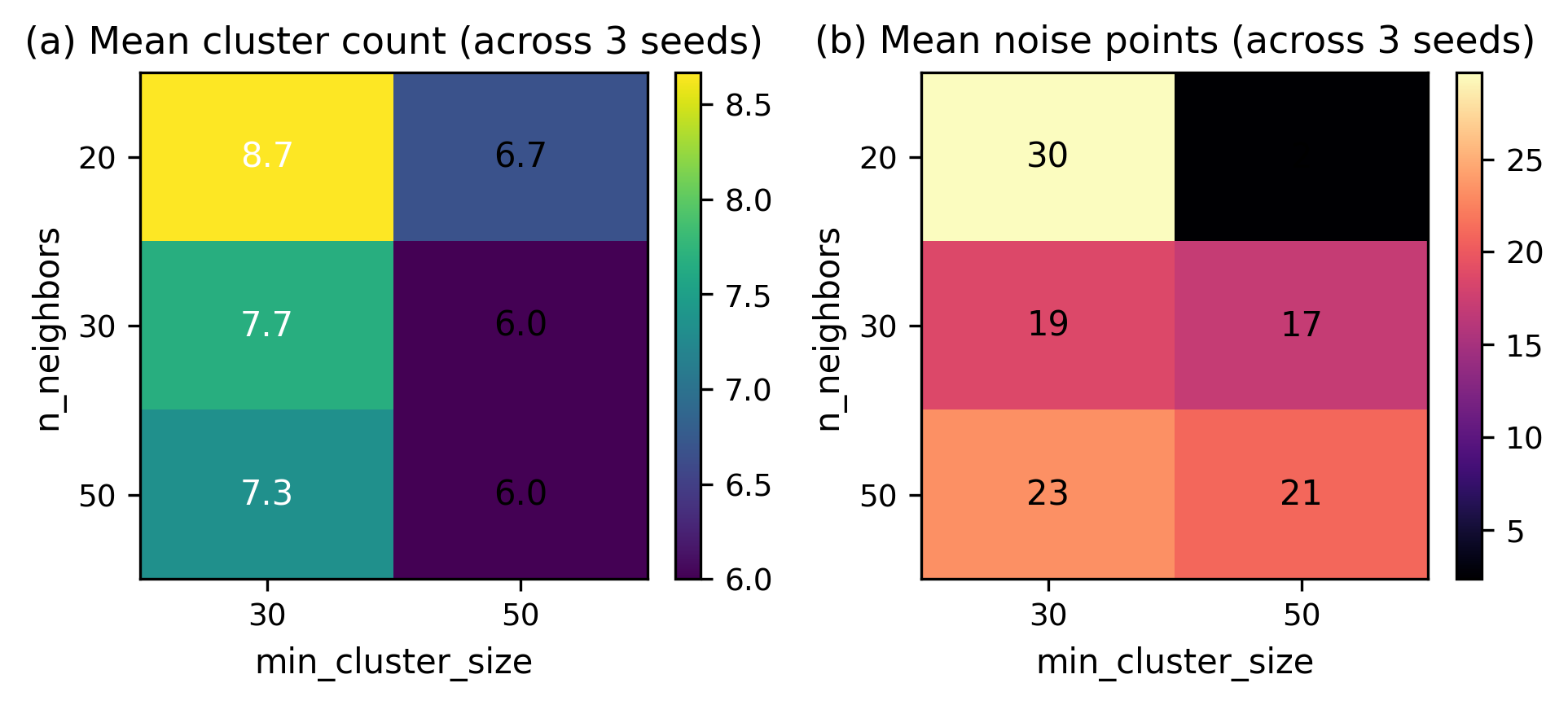

**Fig. S11.** **UMAP and HDBSCAN stability across seeds and parameter settings.** (a) Mean cluster count across three random seeds for each combination of n_neighbors and min_cluster_size. (b) Mean number of HDBSCAN noise points. Cluster count ranges from 5 to 9 across the sweep, with the headline value of 7 falling near the centre of the distribution.

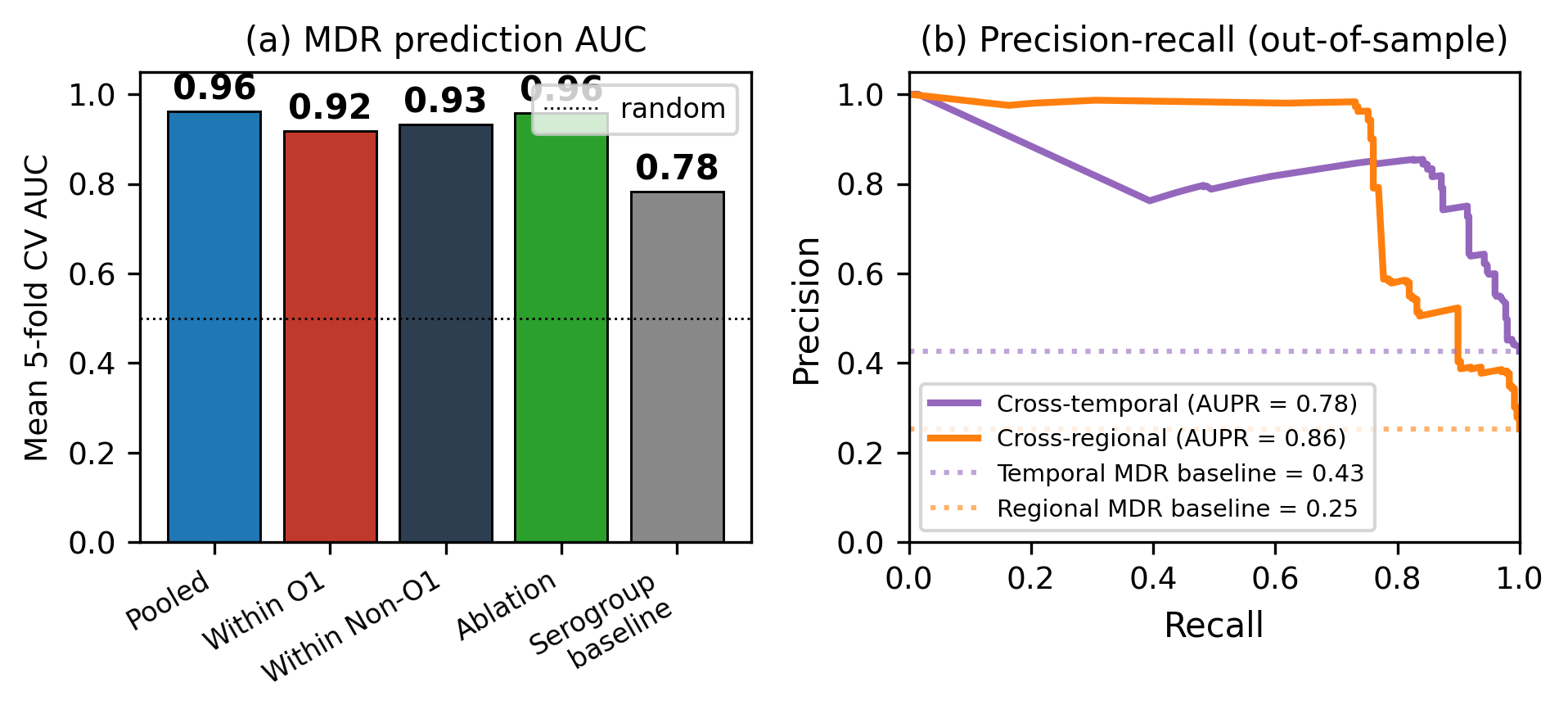

**Fig. S12.** **MDR classifier performance across analysis settings.** (a) Mean 5-fold cross-validated AUC under five settings: pooled, within O1 only, within Non-O1 only, pooled with the six most-diagnostic defence types removed, and a serogroup-only logistic baseline. (b) Precision-recall curves under the cross-temporal (train <=2005, test >2005) and cross-regional (train South Asia plus Africa, test elsewhere) protocols, with the test-set positive-rate baseline shown as a dotted line for each split.
